# A slc38a8 mouse model of FHONDA syndrome faithfully recapitulates the visual deficits of albinism without pigmentation defects

**DOI:** 10.1101/2023.08.19.553949

**Authors:** Ana Guardia, Almudena Fernández, Davide Seruggia, Virginie Chotard, Carla Sánchez-Castillo, Oksana Kutsyr, Xavier Sánchez-Sáez, Esther Zurita, Marta Cantero, Alexandra Rebsam, Nicolás Cuenca, Lluís Montoliu

**Affiliations:** Department of Molecular and Cellular Biology, National Centre for Biotechnology (CNB-CSIC), 28049 Madrid, Spain; Centre for Biomedical Network Research on Rare Diseases (CIBERER-ISCIII), 28029 Madrid, Spain; Sorbonne Université, INSERM, CNRS, Institut de la Vision, F-75012 Paris, France; Department of Physiology, Genetics and Microbiology, University of Alicante, San Vicente del Raspeig Road W/N, 03690, Alicante, Spain; Department of Optics, Pharmacology and Anatomy, University of Alicante, San Vicente del Raspeig Road W/N, 03690, Alicante, Spain; St. Anna Children’s Cancer Research Institute (CCRI), Vienna, Austria, and CeMM Research Center for Molecular Medicine of the Austrian Academy of Sciences, Vienna, Austria

## Abstract

**Purpose:** We aimed to generate and phenotype a mouse model of FHONDA (Foveal Hypoplasia, Optic Nerve Decussation Defects, and Anterior Segment Dysgenesis), a rare disease associated with mutations in *SLC38A8* that causes severe visual alterations similar to albinism without affecting pigmentation.

**Methods:** The FHONDA mouse model was generated with CRISPR (Clustered Regularly Interspaced Short Palindromic Repeats)/Cas9 technology using an RNA guide targeting the *Scl38a8* murine locus. The resulting mice were backcrossed to C57BL/6J. Melanin content was measured using spectrophotometry. Retinal cell architecture was analyzed through light and electron microscopy. Retinal projections to the brain were evaluated with anterograde labelling in embryos and adults. Visual function was assessed by electroretinography (ERG) and the optomotor test (OT).

**Results:** From numerous *Slc38a8* mouse mutant alleles generated, we selected one that encodes a truncated protein (p.196Pro*, equivalent to p.199Pro* in the human protein) closely resembling a mutant allele described in patients (p.200Gln*). *Slc38a8* mutant mice exhibit wild-type eye and coat pigmentation with comparable melanin contents. Subcellular abnormalities were observed in retinal pigment epithelium cells of *Slc38a8* mutant mice. Anterograde labelling experiments of retinal projections in embryos and adults showed a reduction of ipsilateral fibers. Functional visual analyses revealed a decreased ERG response in scotopic conditions and a reduction of visual acuity in mutant mice measured by OT.

**Conclusions:** *Slc38a8* mutant mice recapitulate the phenotype of FHONDA patients concerning their normal pigmentation and their abnormal visual system, as observed in all types of albinism. These mice will be helpful in better understanding the pathophysiology of this genetic condition.

## Introduction

Albinism is a rare genetic condition affecting 1 in 10,000 to 20,000 people in European and Western populations. It is caused by mutations in at least 22 genes, 21 of which have already been identified^1–4^. All types of albinism result in severe visual impairments, with variable degree of hypopigmentation. Visual abnormalities commonly found in patients with albinism include foveal hypoplasia, iris transillumination and misrouted connections between retinal ganglion cells and visual brain nuclei. Consequently, individuals with albinism experience reduced visual acuity, photophobia, nystagmus and limited stereoscopic vision^5^.

In 2013, a novel syndrome called FHONDA (Foveal Hypoplasia, Optic Nerve Decussation Defects, and Anterior Segment Dysgenesis) was reported as a new recessive rare inherited disorder^6^. The visual impairments observed in FHONDA patients were similar to those of albinism, with the exception of anterior segment dysgenesis, only detected in some cases^6–9^. Notably, FHONDA patients have no pigmentation defects, meaning they have normal skin, hair, and eye color^6,7,10,11^. Subsequently, mutations in the *SLC38A8* gene were linked to FHONDA^7^. *SLC38A8* is expressed in the human eye and brain and encodes a transmembrane neuron-specific amino acid transporter known as SNAT8 that transports L-glutamine, L-alanine, L-arginine, L-histidine and L-aspartate using a Na+-dependent mechanism^12^.

Following the discovery that mutations in the *SLC38A8* gene caused FHONDA in a limited number of families^7^, additional patients with new mutations in this locus were reported^8,13–19^. Despite the increasing cohort of patients and mutations, our understanding of the underlying mechanism of FHONDA pathogenesis has not improved significantly.

Animal models, primarily mice, have been instrumental to study the etiology and pathophysiology of albinism, particularly in the pigmentary and visual systems. The majority of mouse models of albinism have been developed by targeting *Tyr*, homologous to the human *TYR* locus, whose mutations cause oculocutaneous albinism type 1 (OCA1)^3,20–22^. These mouse models have been crucial, for example, in uncovering the significant role of L-DOPA in the visual impairments related to albinism^23^.

More recently, mouse models have also contributed to our understanding of the retinal alterations seen in the latest type of albinism, OCA8, caused by mutations in the *DCT* gene^24,25^. However, no animal model has been made available for investigating FHONDA, apart from some morpholino knockdown experiments conducted in Medaka fish, targeting both *Slc38a8* orthologues in the fish, and resulting in microphthalmia, lens defects and fissure coloboma, with varying penetrance, and without any pigmentation defects, as in human FHONDA patients^7^.

Here, we present the generation and phenotypic analysis of the first mouse model of FHONDA. Our aim was to replicate one of the first mutations reported in FHONDA patients (p.Gln200*) by targeting the murine *Slc38a8* locus with CRISPR/Cas9 tools. We obtained several mutant alleles and focused on one (p.Pro196* in mice, equivalent to p.Pro199* in human) that closely mimicked the truncated protein found in FHONDA patients. The resulting *Slc38a8* genome-edited mice exhibited a missing carboxy-end of the protein but maintained wild-type pigmentation levels in the eyes and coat, just like FHONDA patients.

Light and electron microscopy analyses revealed subcellular alterations in the retinal pigment epithelium cells of *Slc38a8* mutant mice. Anterograde labelling of retinal fibers in embryos and adult mice showed a reduced uncrossed projection, consistent with other mouse models of albinism.

Finally, we investigated the visual function of these mice with electroretinography (ERG) and optomotor test (OT). ERG analysis revealed differences in scotopic conditions. The OT demonstrated a statistically significant reduction in visual acuity in *Slc38a8* mutant mice.

This FHONDA mouse model will play a crucial role in further analyses aimed at understanding the underlying molecular mechanisms behind the visual defects in this rare type of albinism, while preserving pigmentation.

## Methods

### Mice

All experimental procedures involving mice were validated by the local CNB Ethics committee on Animal Experimentation. These procedures were then favorably evaluated by the institutional CSIC Ethics Committee and approved by the Autonomous Government of Madrid, in accordance with Spanish and European legislation. All mice were housed at the registered CNB animal facility, where they had ad libitum access to food (regular rodent chow) and water. They were maintained on a light/dark cycle of 08:00–20:00. The authors strictly adhere to the ARVO Statement for the Use of Animals in Ophthalmic and Vision Research. Both male and female mice were used indistinctly in the experiments.

### CRISPR/Cas9 genome editing in mouse fertilized eggs

CRISPR-Cas9 reagents were prepared as described^21,26,27^. Briefly, a synthetic guide RNA (sgRNA) sequence (5’-CAATACTACCTGTGGCCCC-3’), targeting the exon 4 of the *Slc38a8* gene was obtained by *in vitro* transcription. A solution of Cas9 mRNA (25 ng/μl) and the sgRNA (60 ng/μl) was microinjected into 112 mouse B6CBAF2 (Harlan) fertilized oocytes. Subsequently, the 55 surviving embryos (49%) were transferred to three foster females, resulting in the birth of 27 (49%) newborn mice. To confirm successful genome editing, all mice born were screened using the T7 Endonuclease I assay and PCR with these two primers: Fw 5’-AAAGAGATATCCCAGTTGCCCTC-3’ and Rv 5’-ATGGCAAAACCAACAGTCTTCAG-3’. As a result, 24 (89%) founder mosaic individuals were identified.

To deconvolute multiple alleles found in each founder, the region surrounding the target site was cloned, and several clones were subjected to DNA sequencing using the Sanger method resulting in the identification of 16 different mutant alleles. Founder #A8578, carrying the selected mutation (p.196Pro*, equivalent to human p.199Pro*), was utilized to generate the first *Slc38a8* heterozygous (+/−) and homozygous (−/−) animals. Thereafter, this mutant allele was backcrossed five times to the C57BL/6J (Charles River) genetic background, which we used as wild-type pigmented control, therefore the mutants must be formally called B6.CB-*Slc38a8^emA^*^8578^*^/Lmon^*, according to the recommended mouse strain nomenclature^28^. These mutant mice will soon be made available from the EMMA/Infrafrontier repository. As albino control mice, we used B6(Cg)-*Tyr^c-2J/J^* (The Jackson Laboratory) animals.

### Eye pigmentation measurements

The melanin content in adult eye extracts was measured via spectrophotometry, as previously described^21^. Whole mounted retinas and irises were prepared as outlined before^29^. We used Image J for the quantification of iris pigmentation in +18,5 dpc (days post coitum) embryos.

### Histological analysis

Mouse optic cups were fixed in modified Davison solution^30^ and horizontal sections (5 μm thick) were prepared following methods previously described^29^ for light microscopy assessment. These sections were counterstained with cresyl violet.

For electron microscopy, mouse optic cups were fixed with 2% paraformaldehyde and 2% glutaraldehyde. Subsequently, they were postfixed with 1% osmic tetroxide in 0,8% potassium ferricyanide for 1 hour at +4°C. After fixation, the retinas were dehydrated and embedded in resin epon 811. From the embedded tissues, semithin sections (0,5 μm) were obtained and thin sections (50-70 nm) were counterstained with uranyl acetate for 30 minutes at room temperature and with lead citrate for 2 min on golden grids. The samples were then observed using a transmission electron microscope (Jeol JEM 1011-100kV). Each genotype and microscopy technique required the use of at least two adult mice, aged between 2 and 3 months old.

### Labeling of retinal pathways

Anterograde labeling of retinal ganglion cell fibers in +18,5 dpc embryos (10-20 per genotype) was carried out using DiI (1,1′-dioctadecyl-3,3,3′,3′-tetramethylindocarbocyanine perchlorate, Invitrogen) neuronal tracer, following the previously described methods^23,31^.

For adults (6 to 8 individuals per genotype, aged 2-3 months old), anterograde labeling of retinal ganglion cell projections was conducted using intraocular injection of the fluorescent neuronal tracers CTB-AlexaFluor 555 and 647 (#C34776 and #C34778, Thermo Fisher, Waltham, Massachusetts), iDISCO+ brain clearing and 3D imaging with lightsheet microscopy (2X acquisition magnified 0,8X, 3 μm step size) and as reported in previous studies^32,33^. Volume quantifications were performed using a virtual reality headset (Oculus Quest 2, Meta, Menlo Park, California) and Syglass software (Syglass, version 1.2.79, Morgantown, West Virginia). After converting the stack of optical slices into an Imaris file (.ims) using ImarisFileConverter (Bitplane, Oxford Instruments, version 9.5.1, Abingdon, UK), quantifications were performed blindly on the Syglass file (.syg), subsampled to 8 bits (voxel 3.777 x 3.777 x 3 µm). We selected only brains where contralateral superior colliculus staining was evenly distributed, testifying of homogeneous injection. Manual segmentation and thresholding were used to isolate voxels corresponding either to the dLGN ipsilateral or contralateral projection. The threshold was chosen according to the edge of the contralateral SC signal. The signal from the contralateral projection of the dLGN, without thresholding, was used to estimate the total volume of the dLGN, used to normalize volume changes that may occur during clearing. We used a custom algorithm, generated by Syglass engineers in Python, to extract the number of voxels (in units) from each of the channels, converted in .csv files for further volume calculations. Figures were created using Adobe Illustrator (Adobe Creative Cloud 2023), and Imaris.

### Electroretinographic recordings

Scotopic flash-induced ERG responses were recorded in both eyes of the mice as described previously^34^. Briefly, following an overnight phase of dark adaptation, the mice were anesthetized (ketamine 100mg/kg and xylazine 5mg/kg) and kept on a thermal blanket at 38°C. The pupils were dilated using 1% tropicamide (Alcon Cusí, Barcelona, Spain) and ophthalmic gel was used to improve electric contact (Viscotears, Novartis, Barcelona, Spain). A platinum needle was inserted under the skin between the eyes as a reference electrode and ground needle electrode was inserted in the base of the tail. All the procedure was performed in a Faraday cage. The DTL fiber recording electrodes (Retina Technologies, Scranton, PA, USA) were placed in both corneas and the retinas were stimulated by light flashes generated by a Ganzfeld stimulator. Seven stimuli with increasing luminance were presented to the animals. To test the cone contribution to the scotopic ERG responses, double flashes with 1 log cd·s/m2 intensity and an interstimulus interval of 1 s were applied. A 10 second interval was left between stimuli in the case of dim flashes (−5 to −0.6 log cd·s/m2), and up to 20 s for the brightest flashes (0 to 1 log cd·s/m2). ERG records were amplified and band-pass filtered (1-1000 Hz, without notch filtering) using a DAM50 data acquisition board (World Precision Instruments, Aston, UK). A PowerLab system (AD Instruments, Oxfordshire, UK) was used for the presentation of stimuli and data acquisition (4 kHz). The a-wave amplitude was measured from the prestimulus baseline to the most negative trough. The b-wave amplitude measurement was taken from the trough of the a-wave to the peak of the b-wave.

### Optomotor test

To assess the visual acuity of the mice, an Argos optomotor system (Instead Technologies, Elche, Spain, was deployed) was used. The mice were awake and unrestrained. The system is composed of a chamber enclosed by four computer screens facing each other, and a central platform for accommodating the mice (as shown in **Figure 8E**). The visual stimuli used were vertically configured gratings that would pivot around the mouse on a horizontal plane for a duration of 5 seconds, possessing an initial spatial frequency of 0.088 cycles/degree and a contrast of 100%. The responses of the mice were captured by a camera stationed at the topmost part of the system. The mice’s smooth head movements, aligned with the direction of the rotating gratings, were scrutinized by an expert observer. The visual acuity threshold was denoted by the spatial frequency that elicited the mouse’s final response.

### Optical Coherence Tomography (OCT)

A Spectralis OCT system (Heidelberg Engineering, Heidelberg, Germany) allowed acquisition of high-resolution optical coherence tomography images of the central retinal region. Mice were anesthetized via an intraperitoneal injection of ketamine (100 mg/kg) and xylazine (5 mg/kg), and then placed on a heating pad set to 38°C. The pupils were dilated using a topical application of 1% tropicamide (Alcon Cusí, Barcelona, Spain). Corneal hydration was maintained by applying a droplet of physiological saline to each eye. Then, a contact lens was placed on top to improve image quality. High-resolution cross-sectional OCT images were generated specifically in the central retinal area, above the optic nerve. The outer nuclear layer (ONL) and the total retinal thickness, (excluding the retinal pigment epithelium (RPE), were measured across the OCT cross-sectional images at ten discrete points, each separated by 200 µm.

### Statistical analysis

The statistical analyses were conducted with GraphPad Prism (Graph Pad Software, San Diego, CA, USA). Depending on the assay, we used 1-factor or 2-factor ANOVA, with Bonferroni correction. The unpaired Kruskal-Wallis test (without assuming homogeneity of variances) allowed non-parametric comparisons of CTB labeled volumes, followed by Dunn’s multiple comparison test.

## Results

To generate a mouse model of FHONDA we employed a CRISPR-Cas9 approach to target and inactivate the *Slc38a8* murine locus. Our goal was to replicate one of the mutant alleles detected in FHONDA patients (p.200Gln*)^7^, located in exon 4. The human SLC38A8 and mouse Slc38a8 proteins share 82% identity. The p.200Gln position in the human protein sequence corresponds to the p.197Gln in mice (**Supplementary Figure 1**). A unique site for CRISPR targeting was found, where the protospacer adjacent motif (PAM) overlapped with the triplet encoding p.197Gln. RNA reagents, encoding the Cas9 nuclease and the specific RNA guide, were transcribed *in vitro* and microinjected into mouse B6CBAF2 fertilized oocytes following established protocols^21,26^.

Of of the 27 founder mice born, at least 24 (89%) showed signs of successful genome editing in the expected region of the *Slc38a8* locus using the T7 endonuclease I assay, PCR and DNA sequencing analyses. We sequenced all mutant alleles found in founder mice and identified at least 16 different alleles (**Figure 1**). Among these, one encoded a truncated protein that closely resembled the desired mutation (p.196Pro*). The new STOP codon appeared in the triplet encoding the previous amino acid. We selected this mutation for subsequent analysis after confirming germline transmission and obtaining heterozygous and homozygous mice carrying the selected *Slc38a8* mutation (**Supplementary Figure 2**).

**Figure 1.**
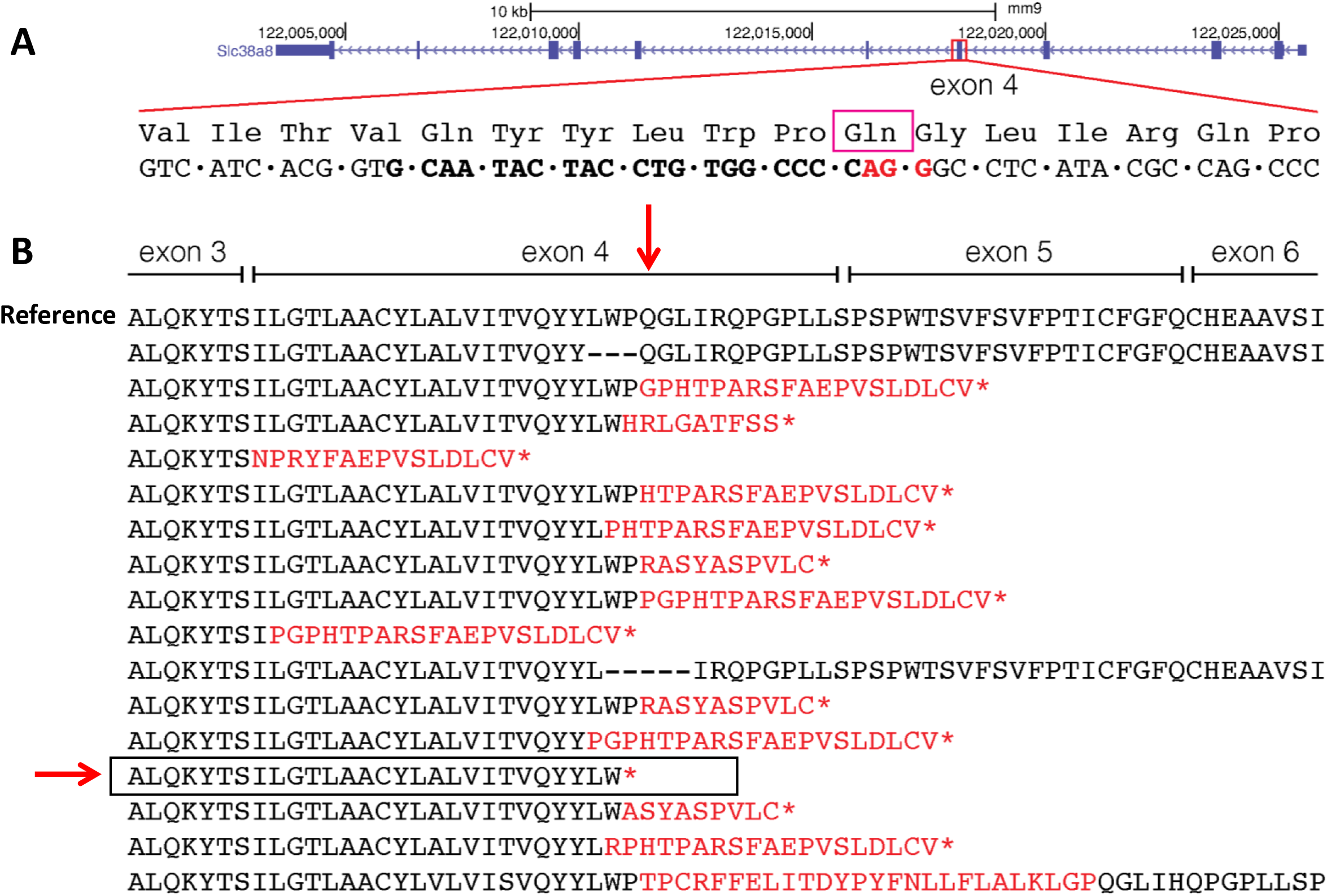
Generation of a *Slc38a8* mouse mutant with CRISPR-Cas9 technology. **A.** Schematic representation of the *Slc38a8* murine locus at chromosome 8, opposite DNA strand (UCSC Genome Browser on Mouse July 2007, NCBI37/mm9). Part of the exon 4 is expanded to show some of the amino acids encoded, including p.197Gln (equivalent to human p.200Gln, indicated by a purple square, see **Supplementary Figure 1**). The DNA sequence underneath contains the sgRNA used (in bold, in black) and the PAM sequence (in bold, in red). **B**. Wild-type Slc38a8 mouse protein sequence (reference) and 16 predicted proteins derived from the 16 mutated alleles identified in CRISPR genome-edited mice. The p.197Gln is indicated by a vertical red arrow. In red, erroneous amino acids generated by the corresponding mutation. An asterisk identifies a newly created STOP codon. The selected FHONDA mutant allele (p.196Pro*), carried by mouse mosaic founder #A8578, is indicated by an horizontal red arrow and framed.

Subsequently, we backcrossed the *Slc38a8* mutant mice five times onto C57BL/6J genetic background and generated the heterozygous (+/−) (**Figure 2C**) and homozygous (−/−) (**Figure 2D**) *Slc38a8* mutants required to perform the rest of phenotypic analyses. The resulting *Slc38a8* (−/−) mice exhibited wild-type pigmentation, undistinguishable from control pigmented mice (**Figure 2A**) and similar to FHONDA patients, in contrast with the albino B6(Cg)-*Tyr^c-2J/J^* mouse that lack pigmentation in both hair and eyes (**Figure 2B**).

**Figure 2.**
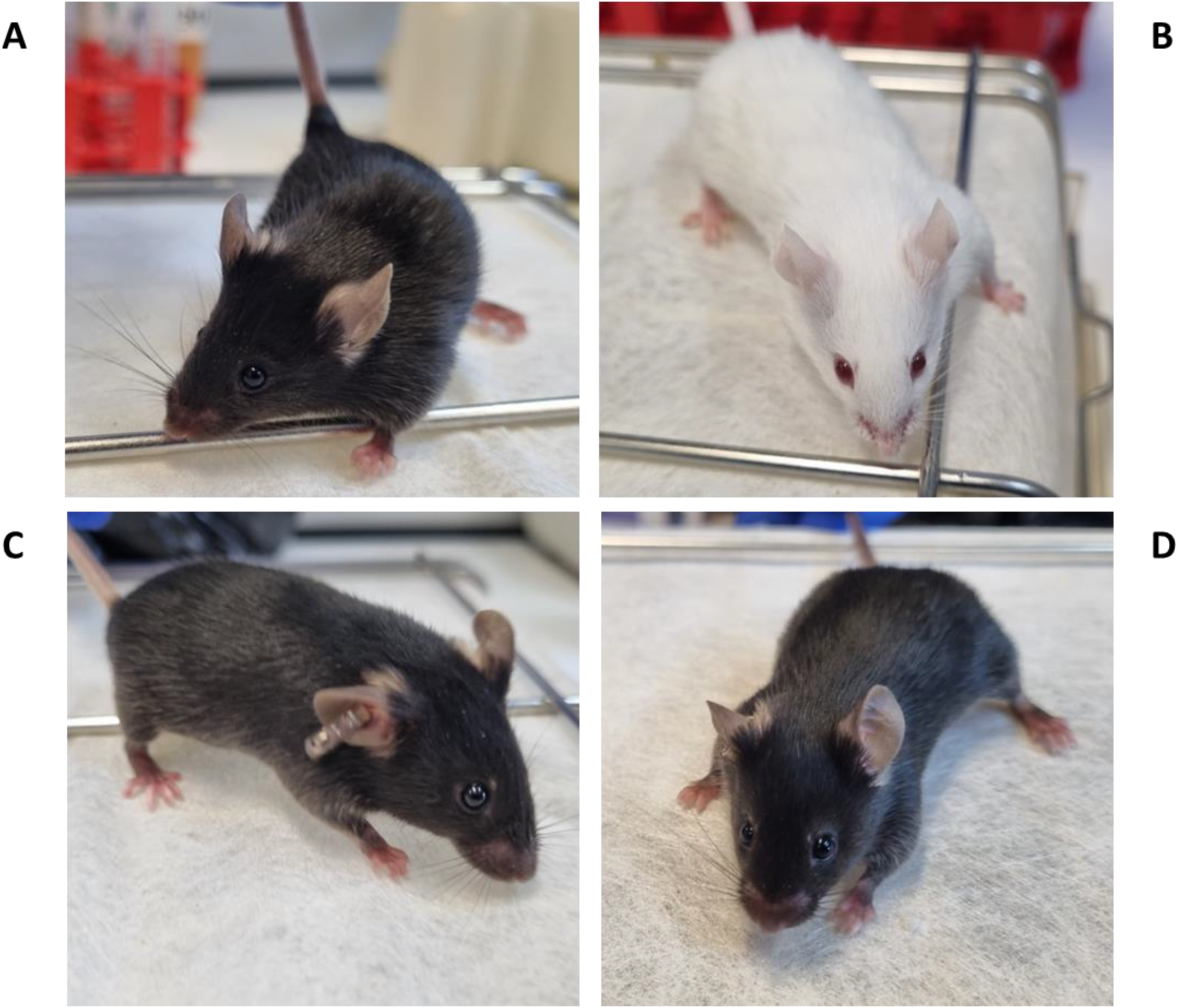
Coat color of *Slc38a8* mutant mice. **A.** wild-type pigmented control C57BL/6J mouse. **B**. albino control B6(Cg)-*Tyr^c-2J/J^* mouse. **C**. *Slc38a8* heterozygous (+/−) mouse in C57BL/6J genetic background. **D**. *Slc38a8* mutant homozygous (−/−) mouse in C57BL/6J genetic background.

We measured the melanin contents in the eyes of *Slc38a8* heterozygous (+/−) and homozygous (−/−) adult mutants, comparing them with wild-type pigmented C57BL/6J and albino B6(Cg)-*Tyr^c-2J/J^* mutant mice (**Figure 3A**). Our findings showed no statistically significant differences between heterozygous or homozygous *Slc38a8* mutants (**Figure 3B**). Similarly, we observed no pigmentation differences in the iris or whole-mounted retina of these *Slc38a8* mutants when compared with C57BL/6J pigmented animals (**Figure 3C**). Surprisingly, statistically significant differences were found between the iris pigmentation in +18,5 dpc embryos of C57BL/6J pigmented mice and heterozygous *Slc38a8* (+/−) mice.

**Figure 3.**
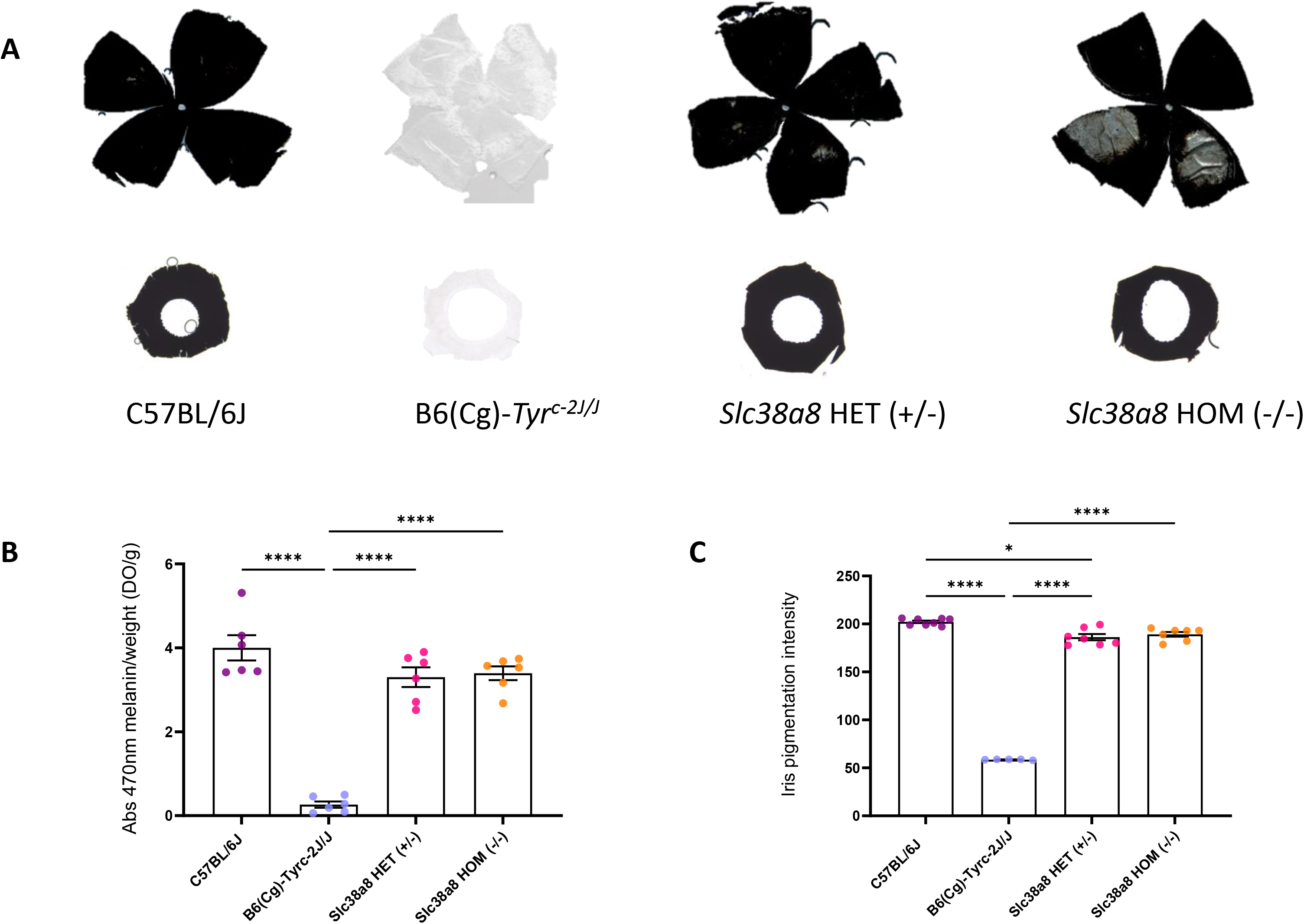
Eye pigmentation of *Slc38a8* mutant mice. **A.** Representative whole-mounted retinas (above) and irises (below) from adult (2-3 months old) wild-type pigmented C57BL/6J, albino B6(Cg)-*Tyr^c-2J/J^*, *Slc38a8* heterozygous “HET” (+/−) and *Slc38a8* homozygous mutant “HOM” (−/−) mice. **B**. Melanin content in the eyes of adult mice from the four genotypes (spectrophotometry). Mean ± SEM, N=6, 1-factor ANOVA with Bonferroni correction for multiple comparisons: * p<0.05, ** p<0.01, *** p<0.005, **** p<0.0001. **C**. Iris pigmentation intensity (ImageJ) in +18,5 dpc embryos. Mean ± SEM, N=5-8, 1-factor ANOVA with Bonferroni correction for multiple comparisons: * p<0.05, ** p<0.01, *** p<0.005, **** p<0.0001.

Next, we evaluated the cellular architecture of the retinas of *Slc38a8* heterozygous and homozygous mutants using light (**Figure 4**) and electron microscopy (**Figure 5**), comparing them to wild-type pigmented C57BL/6J mice and albino B6(Cg)-*Tyr^c-2J/J^*mutants. The retinas of *Slc38a8* heterozygous and homozygous mutants firstly appeared comparable to pigmented retinas under light microscopy (**Figure 4**). However, some empty areas were present in the retinal pigment epithelium (RPE) cells of *Slc38a8* heterozygous and homozygous mutants, particularly in the homozygous mutants (**Figure 4**, arrows). Under electron microscopy, these areas appeared as intracellular vacuoles (**Figure 5**, asterisks), more prominent in the homozygous mutants compared to heterozygous ones, but absent in wild-type or albino RPE cells.

**Figure 4.**
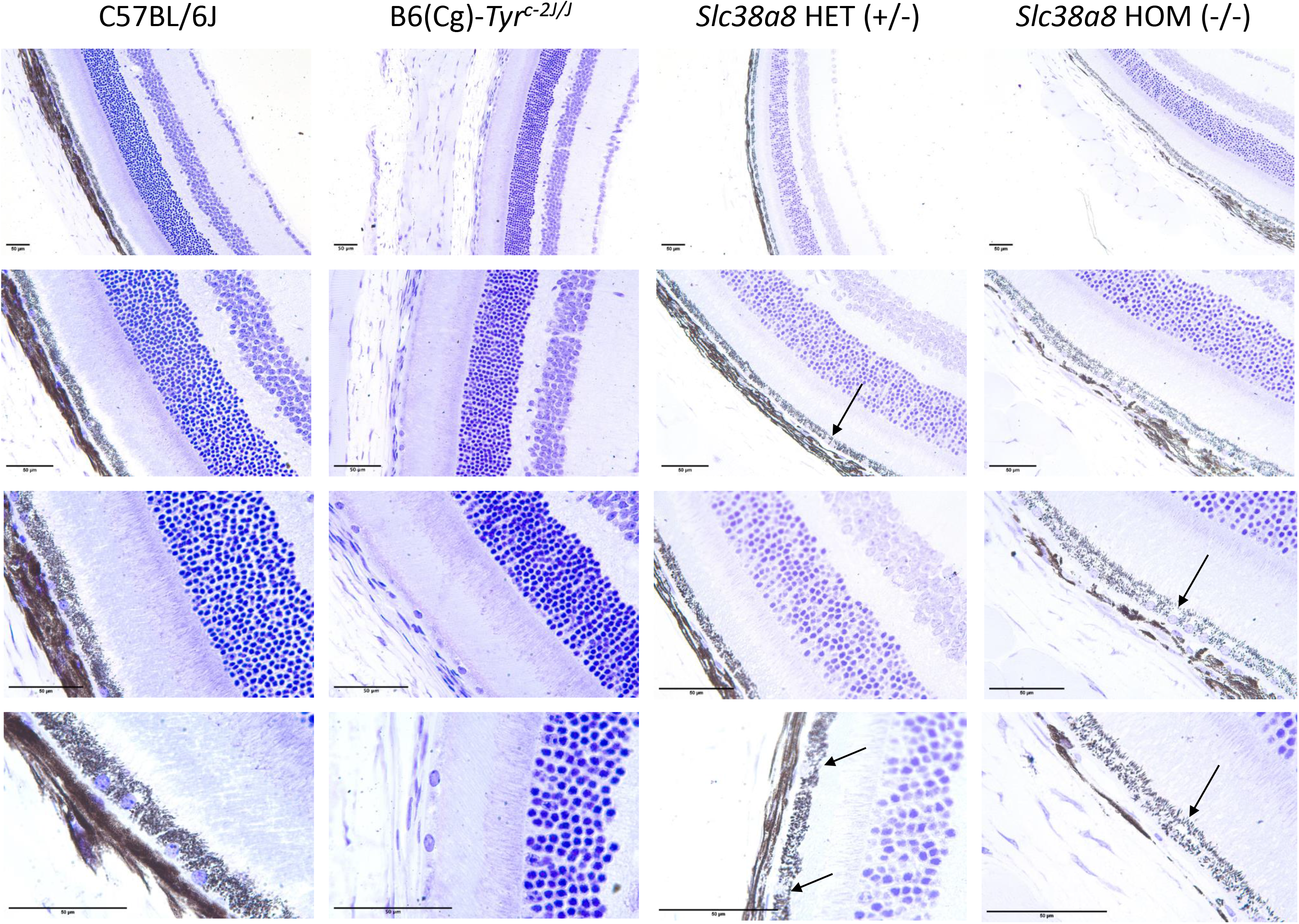
Histological analysis of *Slc38a8* mutant mouse retinas at the light microscope. Representative images of horizontal mouse retina sections (5 μm) counterstained with cresyl violet, from adult (2-3 months old) wild-type pigmented C57BL/6J, albino B6(Cg)-*Tyr^c-2J/J^*, *Slc38a8* heterozygous “HET” (+/−) and *Slc38a8* homozygous mutant “HOM” (−/−). Four different magnifications are shown for each genotype. “Empty” areas within RPE cells in FHONDA mouse retinas are indicated with arrows. The indicated scale bar in all images represents 50 μm.

**Figure 5.**
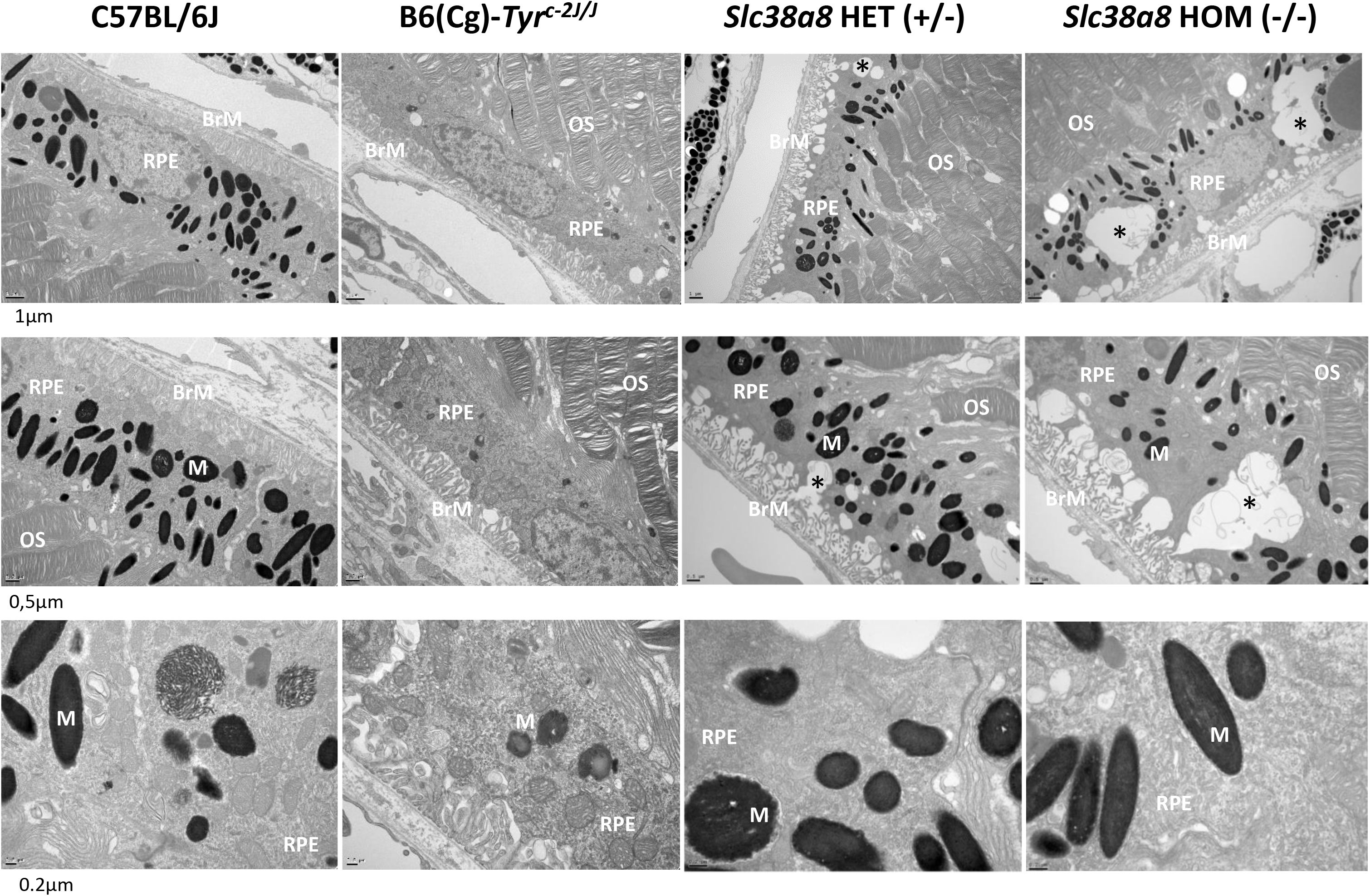
Histological analysis of of *Slc38a8* mutant mouse retinas at the electron microscope. Representative images of mouse retina sections, processed for electron microscopy, from adult (2-3 months old) wild-type pigmented C57BL/6J, albino B6(Cg)-*Tyr^c-2J/J^*, *Slc38a8* heterozygous “HET” (+/−) and *Slc38a8* homozygous mutant “HOM” (−/−). Three different magnifications are shown for each genotype. Subcellular alterations/abnormal vacuoles within RPE cells are indicated with asterisks. The indicated scale bar in all images represents 1 μm for the top row, 0.5 μm for the middle row, and 0.2 μm for the bottom row. RPE, retinal pigment epithelium cells; OS, outer segments of photoreceptor cells; BrM, Bruch’s membrane; M, melanosomes.

Additionally, we measured the number of pigmented melanosomes per RPE cell on electron microscope images, and found no differences between pigmented C57BL/6J, heterozygous and homozygous *Slc38a8* mutants (**Supplementary Figure 3**).

In albino mammals, the projections of retinal ganglion cells to visual nuclei of the brain are affected, with a reduced amount of ipsilateral fibers departing from the optic chiasm^23,35^. We decided to explore these projections in *Slc38a8* mutant mice using two anterograde labeling approaches, in embryos and adults, following established procedures^23,36^.

Unilateral anterograde labeling with DiI (1,1′-dioctadecyl-3,3,3′,3′-tetramethylindocarbocyanine perchlorate) in +18.5 dpc embryos revealed a significant reduction in ipsilateral fibers in *Slc38a8* homozygous mutant mice (**Figure 6A-D**). Indeed, the vast majority showed faint ipsilateral projections (4 out of 9 homozygous mutant embryos) and none of them had an ipsilateral projection as wide and visible as in wild-type mice. These percentages were similar to the reduction observed in B6(Cg)-*Tyr^c-2J/J^*albino control mice, where 8 out of 11 embryos showed faint ipsilateral projections. In contrast, marked ipsilateral projections were observed in 10 out of 12 *Slc38a8* heterozygous embryos, comparable to wild-type pigmented animals (marked projections in 9 out of 12 fetuses, **Figure 6E**).

**Figure 6.**
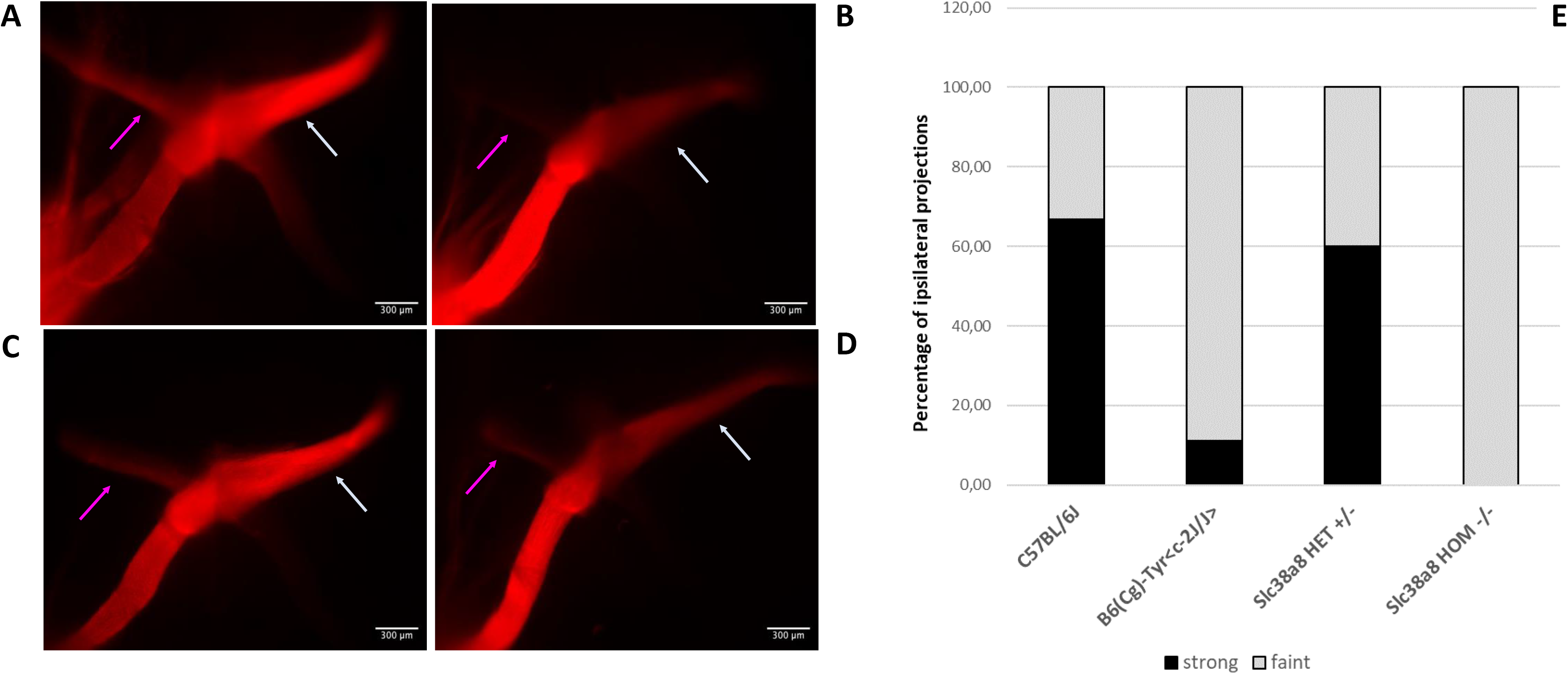
Analysis of retinal ganglion cell projections of *Slc38a8* mutant mice. Unilateral (left eye) anterograde labeling of +18.5 dpc embryos with neuronal tracer DiI. **A**. Representative images of whole-mounted optic chiasms from wild-type pigmented C57BL/6J, albino B6(Cg)-Tyrc-2J/J, *Slc38a8* heterozygous “HET” (+/−) and *Slc38a8* homozygous mutant “HOM” (−/−) mice. Ipsilateral projections are indicated with a purple arrow. Contralateral projections are indicated with a white arrow. **B**. Quantification of ipsilateral projections. Bars are expressed in percentages considering faint (barely visible, shown in grey) or strong (clearly visible, shown in black) uncrossed fibers. Data are derived from N=12 (pigmented), N=11 (albino), N=12 (*Slc38a8* +/−) and N=9 (*Slc38a8* −/−) number of fetuses analyzed. Ipsilateral projections were detected only in N=9 (pigmented, 6 strong and 3 faint), N=9 (albino, 1 strong and 8 faint), N=10 (*Slc38a8* +/−, 6 strong and 4 faint) and N=4 (*Slc38a8* −/−, 0 strong and 4 faint).

We conducted similar anterograde tracing experiment in adult *Slc38a8* heterozygous and homozygous mutant mice (2-3 months old) using the neuronal tracer CTB (Cholera Toxin B-subunit) associated with AlexaFluor 555 (represented in magenta) or AlexaFluor 647 (represented in green) fluorochromes into each eye, respectively (**Figure 7A**) to visualize the entire 3D retinal projections for both eyes using clearing and lightsheet microscopy (**Figure 7B**). We used control wild-type pigmented C57BL6/J, heterozygous S*lc38a8* and homozygous *Slc38a8* mutant mice and observed a reduced ipsilateral projection in homozygous *Slc38a8* mutants, especially in the superior colliculus (SC) when looking at the entire 3D projection (**Figure 7C**) and in the dLGN (**Figure 7D**), which aligns with our results in embryos (**Figure 6**). Indeed, there was a statistically significant reduction in the volume of the ipsilateral projection in the dLGN of homozygous *Slc38a8* mutants compared to C57BL6/J mice in both raw (**Figure 7F**) or normalized volume over the total dLGN volume (**Figure 7G**). To note, the dLGN is smaller in homozygous *Slc38a8* mutants compared to heterozygous *Slc38a8* mice (**Figure 7H**). Overall, the ipsilateral projection is significantly reduced in homozygous *Slc38a8* mutants compared to control wild-type C57BL6/J mice. Furthermore, we observed an ectopic patch in the contralateral dLGN projection of homozygous *Slc38a8* mutants, not present in the heterozygous S*lc38a8* and C57BL6/J wild-type animals (**Figure 7E**) but similar to the ectopic patch described in albino B6(Cg)-*Tyr^c-2J/J^* mutant mice^36^.

**Figure 7.**
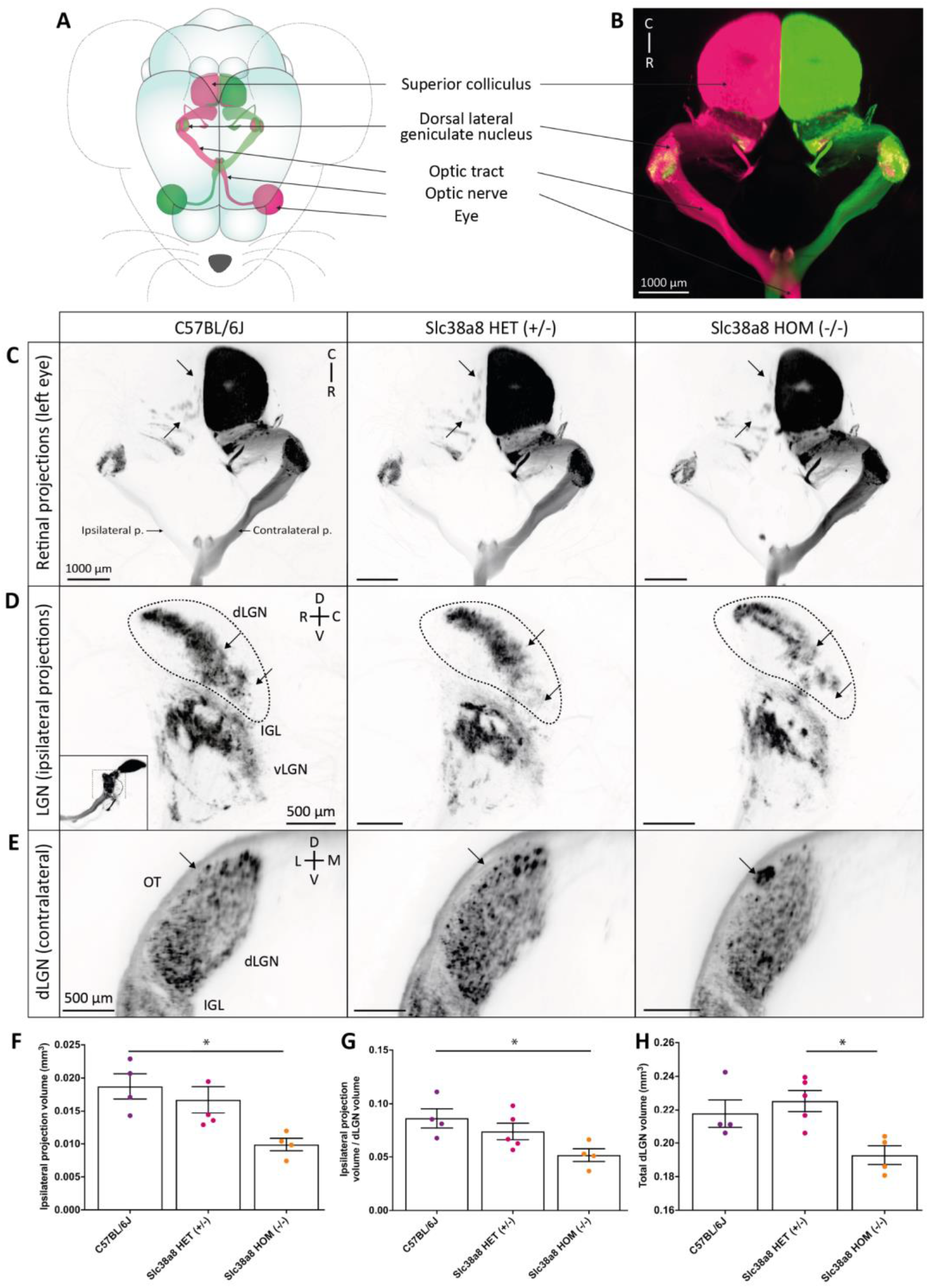
Slc38a8 homozygous mutation results in a major deficit of retinal axonal projections in mice. **A.** Schematic view of retinal projections in different visual targets after anterograde axonal tracing following intraocular injection of Alexa Fluor 555 or 647 conjugated cholera β-subunit toxin (CTB) in each eye. **B**. 3D image of the light sheet acquisition showing structures schematized in (A). Ipsilateral and contralateral projections are well segregated in the dorsal lateral geniculate nucleus (dLGN) of a control C57BL6/J mouse. Ipsilateral projections in the superior colliculus (SC) are not visible due to the intensity of the contralateral staining. **C**. Top view of the entire ipsi- and contralateral retinal projections from one eye in C57BL6/J, Slc38a8 HET (+/−), and Slc38a8 HOM (−/−) mice. The number of ipsilateral projection patches in the SC is notably decreased Slc38a8 HOM (−/−) mice, compared to Slc38a8 HET (+/−) and C57BL6/J. **D**. Lateral view of the retinogeniculate ipsilateral projections in the dLGN, the intergeniculate lamina (IGL) and the ventral LGN (vLGN). The thumbnail image indicates the orientation of the capture, as the LGN was visually isolated from the rest of the projections for clarity’s sake. Arrows indicate areas in the dLGN where major changes were visible in Slc38a8 HOM (−/−) mice compared with Slc38a8 HET (+/−) and C57BL6 mice, including a decrease in projection density. **E**. Optical sections of the caudal dLGN showing an ectopic patch of contralateral projections (arrow) in the Slc38a8 HOM (−/−) mutants at the periphery of the dLGN, adjacent to the optic tract (OT), that was absent in the other genotypes. **F, G**. Volume of ipsilateral projections in the dLGN in C57BL6/J, Slc38a8 HET (+/−), and Slc38a8 HOM (−/−) mice, as raw values (F) or normalized over the volume of the total dLGN (G) show a reduction in the volume of the ipsilateral projections in Slc38a8 HOM (−/−) compared to C57BL6/J mice. **H**. Total dLGN volume used for normalization. Abbreviations: p. projection; D: dorsal; V: ventral; C: caudal; R: rostral; L: lateral; M: medial. Statistics: Kruskal-Wallis test, followed by Dunn’s multiple comparisons; * p<0.05. Small variations in volume were present due to the clearing process. Image colors have been inverted for better visibility in C-E. Bars indicate means and SEM.

Finally, having confirmed the RPE cell alterations and the misrouting of the ganglion cell ipsilateral projections, we proceeded to functionally assess the visual capacity of these *Slc38a8* mutant mice. This investigation aimed to determine whether the observed morphological alterations in the optical tract led to a significant reduction in visual acuity, as it is commonly observed in FHONDA patients.

The same three genotypes (wild-type control C57BL/6J, *Slc38a8* heterozygous and *Slc38a8* homozygous mutants) at adulthood (2-3 months old) were subjected to electroretinography analyses under scotopic conditions to evaluate their retinal function. The analyses of ERG amplitudes showed statistically significant differences in the responses elicited by the higher intensity flashes between wild-type and *Slc38a8* homozygous mutant mice, being smaller in the latter group, as expected (**Figure 8A-B**). In fact, the maximum mean value of the mixed a-wave amplitude in the control group was reduced by 21% (p<0.01) in the *Slc38a8* homozygous mutants (**Figure 8A-B**). Surprisingly, we also detected additional statistically significant differences between heterozygous and *Slc38a8* homozygous mutants, with the mean maximum a- and b-wave amplitude values of homozygous mutants reduced by 20% (p<0.01) and by 15% (p<0.01) respectively when compared to the *Slc38a8* heterozygous group (**Figure 8A-C**). However, when we studied the cone-driven ERG response by a scotopic double flash protocol, the amplitudes of a- and b-waves showed no differences between the three groups (**Figure 8A, D**).

**Figure 8.**
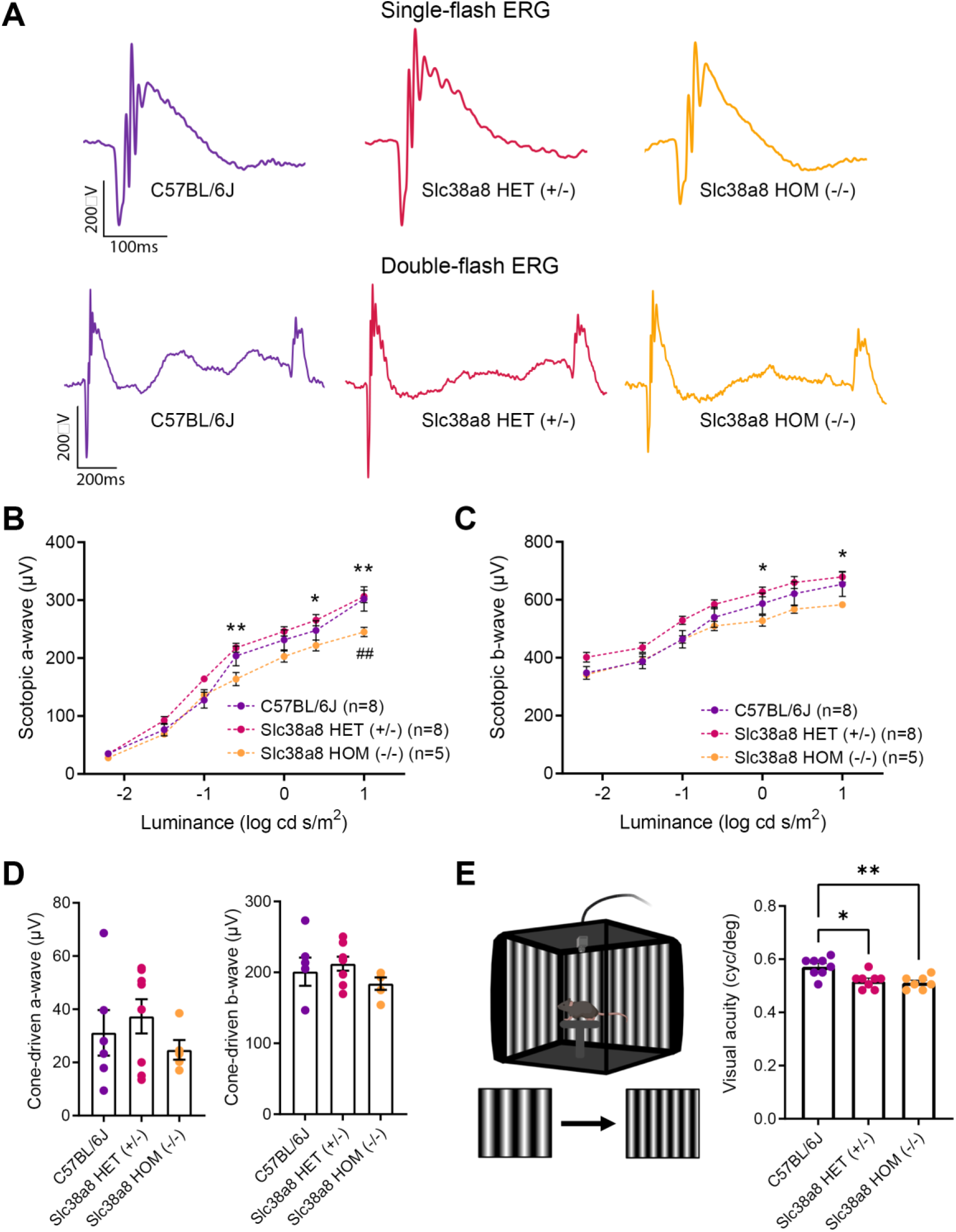
Functional retinal analysis of *Slc38a8* mutant mice. Electroretinogram recordings in scotopic conditions were obtained from adult (2-3 months old) wild-type control pigmented C57BL/6J, *Slc38a8* heterozygous “HET” (+/−) and *Slc38a8* homozygous mutant “HOM” (−/−). **A**. Representative dark-adapted ERG intensity responses to single and double 1 log cd·s/m2 flashes from the three genotypes. **B-C**. Luminance-response curves of the three experimental groups. Each graph includes measurements from scotopic mixed responses depicting a-wave (B) and b-wave (C) amplitudes. **D.** Maximal scotopic cone-driven response from the three genotypes showing maximal a- and b-wave amplitudes. No statistically significant differences were found**. E**. Left: configuration of the optomotor system. Image was created using BioRender (https://biorender.com/). Right: visual acuity measured as the spatial frequency threshold in the three experimental groups. Mean ± SEM, N=5-8. Two-way ANOVA with Bonferroni correction (ERG) and Kruskal-Wallis test and Dunn’s post-hoc test (optomotor test). * p<0.05, ** p<0.01, Asterisks (*) refer to comparison between *Slc38a8* heterozygous (+/−) and *Slc38a8* homozygous (−/−) mice. Hash (#) refers to comparisons between control pigmented C57BL/6J and *Slc38a8* homozygous (−/−) mice.

For a comprehensive evaluation of the vision of these mice, we assessed their visual acuity with the optomotor test. As expected, *Slc38a8* homozygous mutant mice exhibited a statistically significant reduction in visual acuity compared to control C57BL/6J mice. Surprisingly, the *Slc38a8* heterozygous mice also showed a similar reduction in visual acuity. Notably, these two experimental groups did not differ from each other (**Figure 8E**). Furthermore, using OCT, we measured both the thickness of the total retina and of the outer nuclear layer (ONL) comprising photoreceptor nuclei. With this approach, we did not find any statistically significant differences in either of these measurements, revealing that there is no apparent loss of photoreceptor and the retina structure is maintained (**Figure 9**).

**Figure 9.**
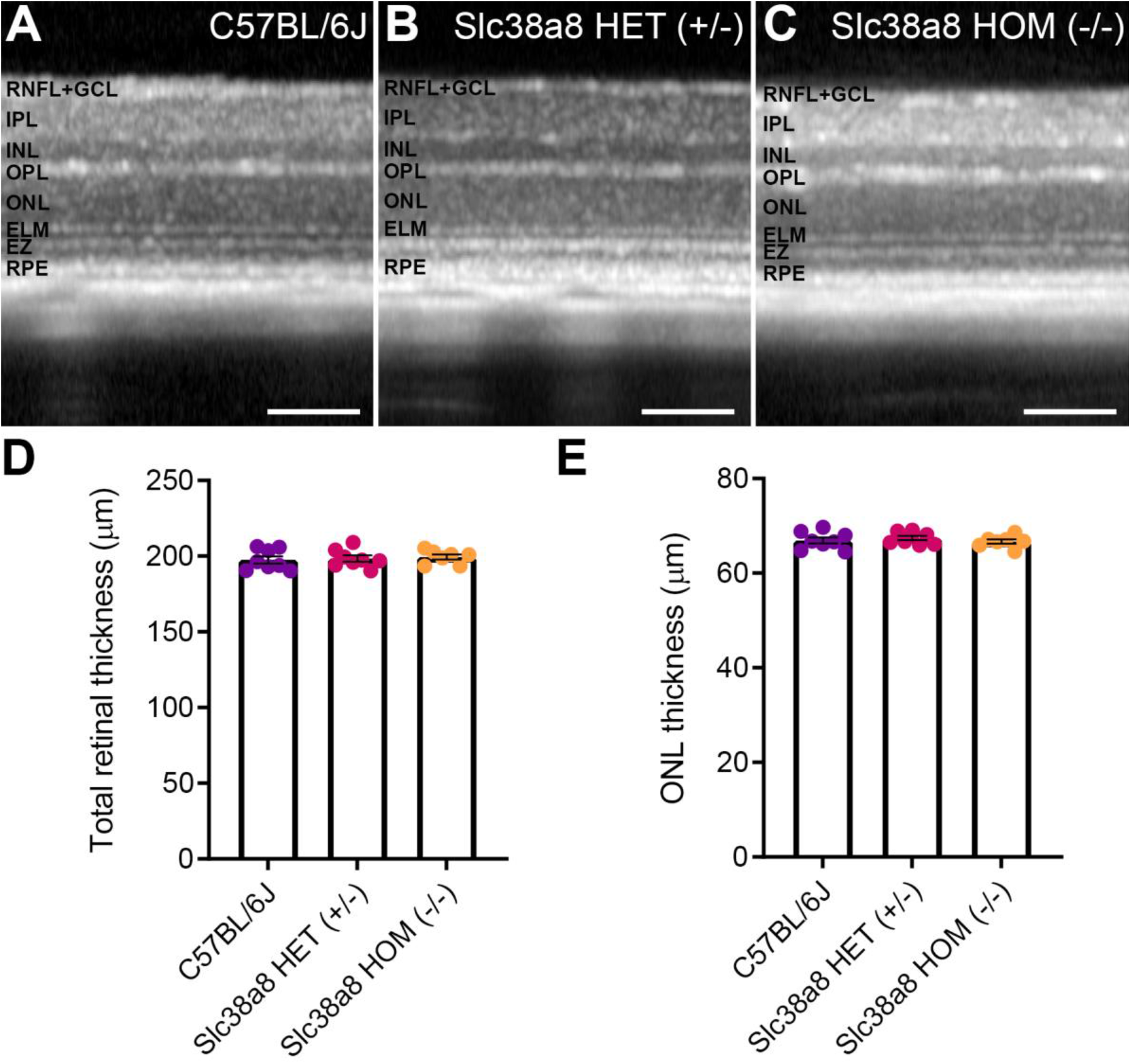
Retinal structure of *Slc38a8* mutant mice evaluated in vivo by optical coherence tomography. Representative OCT images from control pigmented C57BL/6J (A), heterozygous *Slc38a8* (+/−) (B) and homozygous *Slc38a8* (−/−) (C) mutant mice. D Quantification of the total retinal thickness. E. Quantification of the ONL thickness (Outer Nuclear Layer, photoreceptor nuclei). Mean ± SEM, N=5-8. Two-way ANOVA with Bonferroni correction. There are no statistically significant differences. RNFL: retinal nerve fiber layer, GCL: ganglion cell layer, IPL: inner plexiform layer, INL: inner nuclear layer, OPL: outer plexiform layer, ELM: external limiting membrane, EZ: ellipsoids zone, RPE: retinal pigment epithelium. Scale 100 µm.

## Discussion

We generated a mouse model of FHONDA by installing a truncating mutation (p.Pro196*) at the *Slc38a8* locus using CRISPR-Cas9 genome editing tools. This mutant closely resembles one of the first familial mutations reported at the *SCL38A8* locus in FHONDA patients (p.Gln200*)^7^. We subjected this animal model to extensive phenotyping.

Our results indicate that the pigmentation of *Slc38a8* homozygous mutant mice is similar to wild-type pigmented control C57BL/6J mice in terms of coat color, iris pigmentation, melanin content in the eye, mirroring the observation in FHONDA patients that pigmentation is not affected^6,7,11^. However, we noted a statistically significant decrease in pigmentation intensity in the iris of +18,5 dpc *Slc38a8* heterozygous embryos compared with wild-type pigmented C57BL/6J embryos. This slight discrepancy could be related to subtle differences in the genetic background of these animals (98,44% of C57BL/6J genome after five rounds of backcrossing).

Interestingly, we noted subcellular abnormalities during the histological assessment, with empty areas within the RPE cells under light microscopy, identified as large vacuoles inside the RPE cells by electron microscopy. These structures were not present in pigmented C57BL/6J and albino B6(Cg)-*Tyr^c-2J/J^* mice. Vacuolization of RPE cells has been commonly reported in mouse models of other visual pathologies^37^. Besides, the number of pigmented melanosomes within RPE cells of wild-type pigmented C57BL/6J, *Slc38a8* heterozygous and homozygous mutant mice did not exhibit any differences.

We show that the ipsilateral projection is reduced both at the optic chiasm in embryos and in brain targets in adult *Slc38a8* homozygous mutant mice. Reduced ipsilateral ganglion cell projections is a characteristic alteration of the visual tract found in all albino mammals, especially in mouse models of albinism^23,31,35^. Also, Ipsilateral misrouting is a common feature present in all FHONDA patients^9^, although one study documented a FHONDA patient who showed no evidence of chiasmal misrouting^18^. Interestingly, the visual cortex of FHONDA patients receives inputs only from the contralateral eye, while the visual cortex in patients with other types of albinism receives inputs from both eyes, albeit a much smaller proportion from the ipsilateral eye compared to control subjects^38^, suggesting a stronger phenotype following *SLC38A8* mutation.

We evaluated the visual function of *Slc38a8* mutant mice using electrophysiological and behavioral methods. Mixed ERG responses performed under scotopic conditions, where rod photoreceptors are active, revealed statistically significant differences between control C57BL/6J and *Slc38a8* homozygous mutant mice, as well as between *Slc38a8* heterozygous and homozygous mutants. However, none of these differences were observed when we evaluated the cone contribution to the ERG under scotopic conditions. It is worth noting that some FHONDA patients also exhibit normal ERG measurements^19^.

Some of these findings (differences in rod versus cone response) are also consistent with a moderate rod deficit observed in albino mice^23,39^. However, OCT measurements to assess the retinal and ONL thickness did not confirm the reported differences in photoreceptor numbers, but the level of resolution may not be sufficient. Similarly, in FHONDA patients, foveal hypoplasia is not always detected in all individuals^16^.

Behavioral studies with OT clearly demonstrated that *Slc38a8* homozygous mutant mice exhibited a statistically significant reduction in visual acuity, as observed in FHONDA patients^9^. However, the same test detected an unexpected reduction in visual acuity in *Slc38a8* heterozygous animals.

In this study, we generated and phenotyped a FHONDA mouse model that faithfully recapitulates the phenotype observed in FHONDA patients^6–8,13–19,38^, providing valuable insights into the underlying mechanisms and potential therapeutic targets for this condition.

Interestingly, for most tests, *Slc38a8* heterozygous mice behaved similarly to pigmented control animals. Surprisingly, in two tests (subcellular alterations in RPE cells, optomotor test), the same *Slc38a8* heterozygous animals behaved similarly to *Slc38a8* homozygous mutants. These seemingly contradictory results suggest that *Slc38a8* heterozygous mice might also exhibit an altered visual phenotype. It is possible that *Scl38a8* mutations behave in a semi-dominant manner rather than a recessive one, as initially anticipated.

To our knowledge, there has been only one individual described carrying an heterozygous mutation in *SLC38A8* with a partial FHONDA phenotype (nystagmus, but with a normal fovea and normal decussation)^16^. There are published examples of semi-dominant inheritance in rare diseases^40,41^, including recent cases of syndromic albinism^42^. These findings indicate that semi-dominant inheritance is a possibility and should be explored further in FHONDA.

The visual characteristics observed in FHONDA patients, including foveal hypoplasia, nerve misrouting, reduced visual acuity, and nystagmus, closely resemble those seen in other types of albinism and prompted researchers to list FHONDA as a new genetic subtype of albinism where patients do not exhibit any signs of hypopigmentation^4^. The visual phenotype of our FHONDA mouse model, with the morphological, anatomical and functional visual abnormalities detected in *Slc38a8* homozygous mutant mice, further contributes to associate FHONDA with the visual alterations commonly found in albinism.

Similar to FHONDA, ocular albinism type 1 (OA1), associated with mutations in the *GPR143* gene, is also characterized by the visual abnormalities commonly found in albinism with subtle ocular pigmentation defects^43^.

The complex and heterogenous genetic nature of albinism has been a perplexing challenge for researchers over the years. Understanding the range of genes responsible for causing similar retinal phenotypes has only just begun. Recently, researchers have proposed a hierarchical arrangement of all these genes, outlining their relationships based on the phenotypes observed^4^. Interestingly, the proposed ordered list ends with the *GPR143* and *SLC38A8* genes, as the two most downstream effectors of the visual alterations associated with albinism. Our model opens new possibilities to study the mechanisms at the origin of these congenital visual disorders.

## Acknowledgements

This work has been funded by the Spanish Ministry of Economy and Competitiveness under BIO2015-70978-R, the Spanish Ministry of Science and Innovation under RTI2018-101223-B-I00, CIBERER and Fundación Ramón Areces to LM. Additionally, Spanish Ministry of Science and Innovation [FEDER-PID2019-106230RB-I00, 2019] and Generalitat Valenciana IDIFEDER/2017/064, 2017, PROMETEO/2021/024, 2021 supported the work of NC. Funds from INSERM, Sorbonne Université, Retina France and Genespoir supported the work of AR, as well as LabEx LIFESENSES (ANR-10-LABX-65) and IHU FOReSIGHT (ANR-18-IAHU-01) for the Institut de la Vision, a doctoral fellowship from the French Ministry of Education and Research to VC. The authors would like to express their gratitude to C. Lillo (INCyL-USAL, Salamanca, Spain) for help interpreting the electron microscopy images, and to the transgenesis, histology, electron microscopy, advanced light microscopy and mouse embryo cryopreservation facilities at CNB-CSIC for the outstanding support and services provided.

---

**Supplementary Figure 1.**
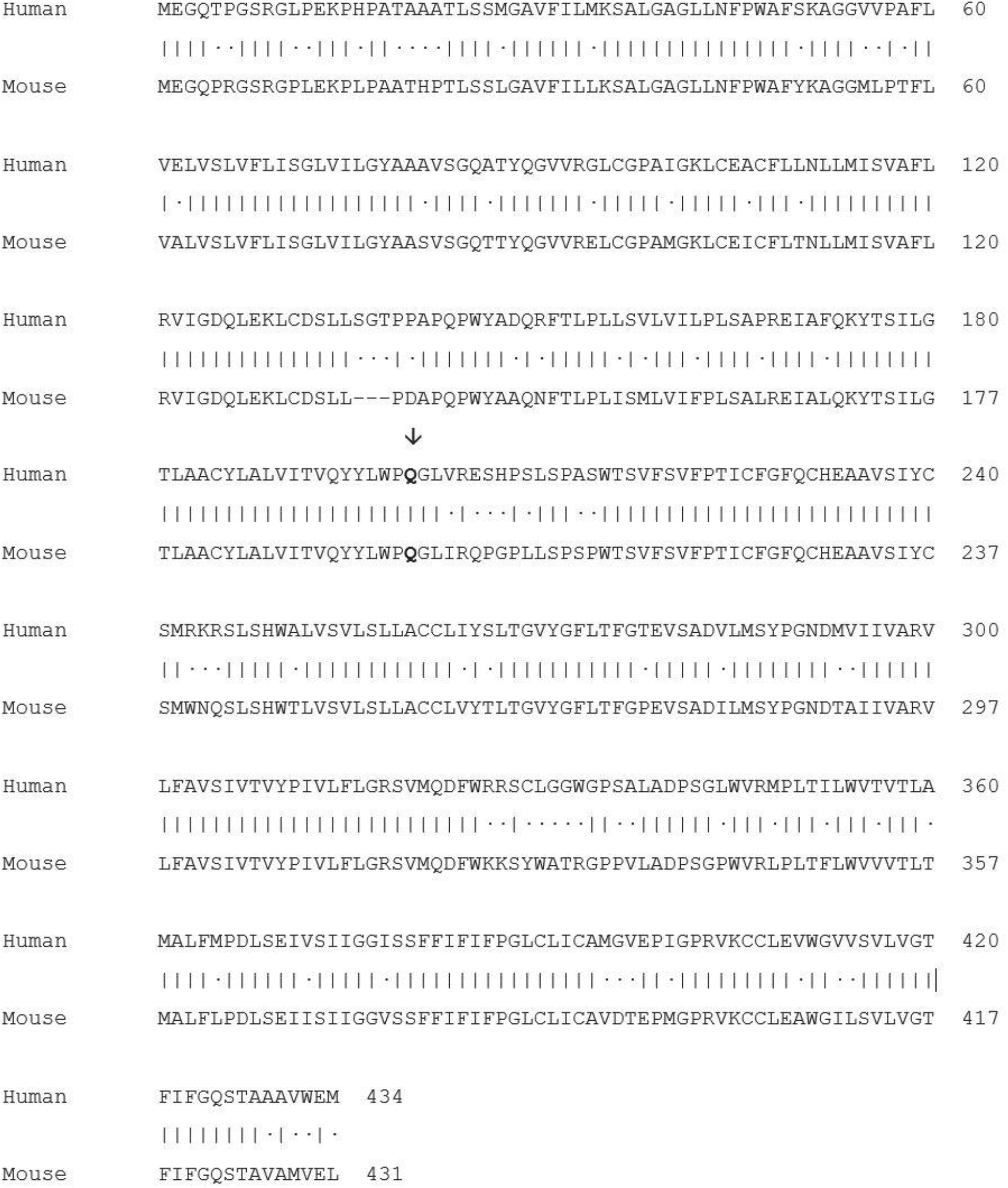
BLASTP (NIH/NCBI) alignment of SLC38A8 human (A6NNN8 UniProtKB, 434 amino acides) and Slc38a8 (Q5HZH7 UniProtKB, 431 amino acids) mouse proteins. The p.200Q (human) and p.197Q (mouse) amino acid residues are highlighted in bold and indicated with a vertical arrow.

**Supplementary Figure 2.**
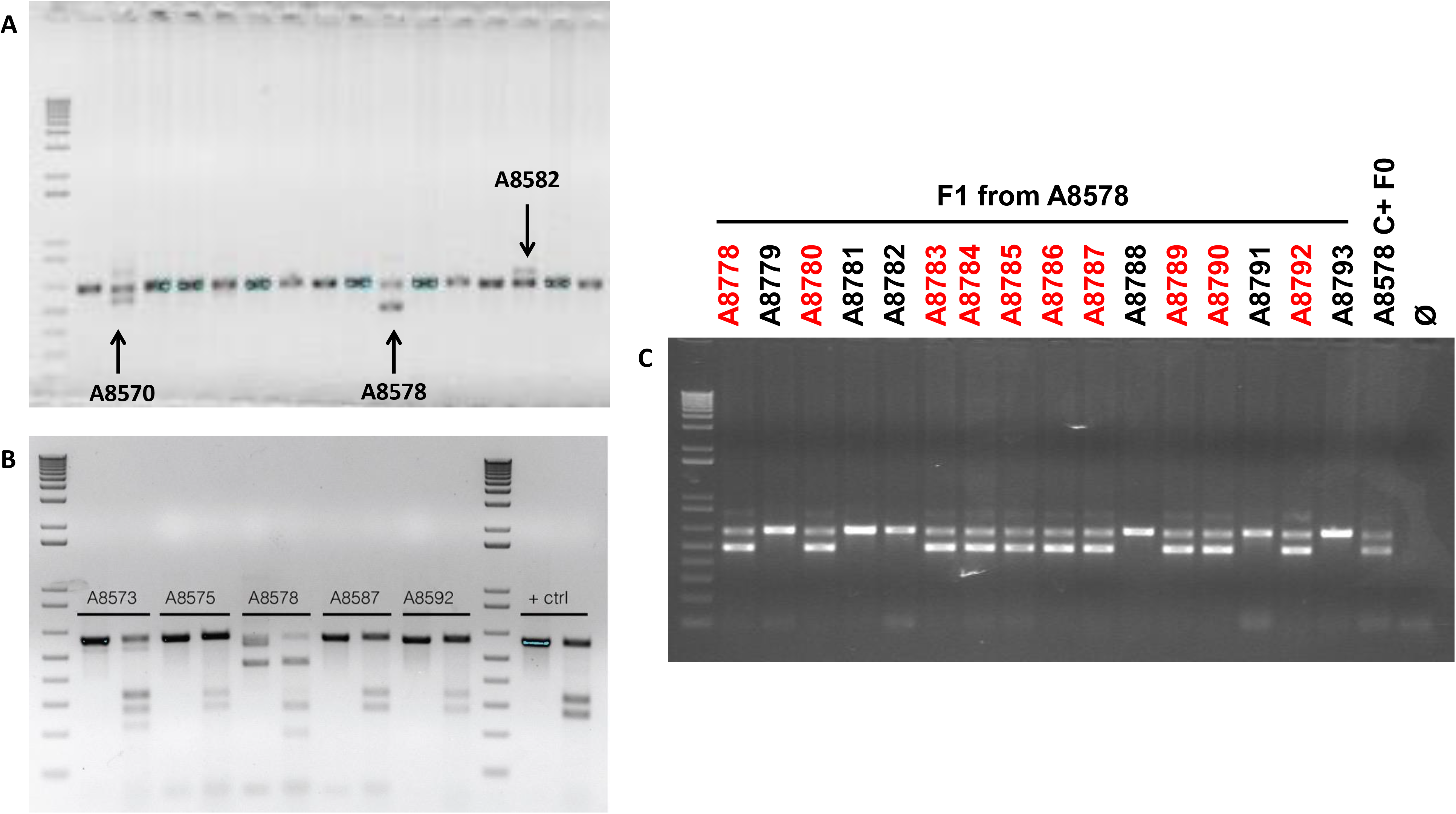
Identification of FHONDA mutant founder (F0) mice after genome editing with CRISPR-Cas9 tools. **A**. Detection of FHONDA mutant founder (F0) mice by PCR. Three positive founder mice are shown, including #A8578, the mouse that carries the p.199Pro* mutant allele eventually selected for this study. Larger bands correspond to insertions and smaller bands correspond to deletions. **B**. Detection of FHONDA mutant founder (F0) mice by T7 Endonuclease I assay, which detects indels (insertions/deletions). Five positive founder mice are shown, including #A8578. **C**. Germline transmission test of FHONDA mouse mutant founder (F0) #A8578. The deleted product (smaller band) is found in numerous F1-derived animals.

**Supplementary Figure 3.**
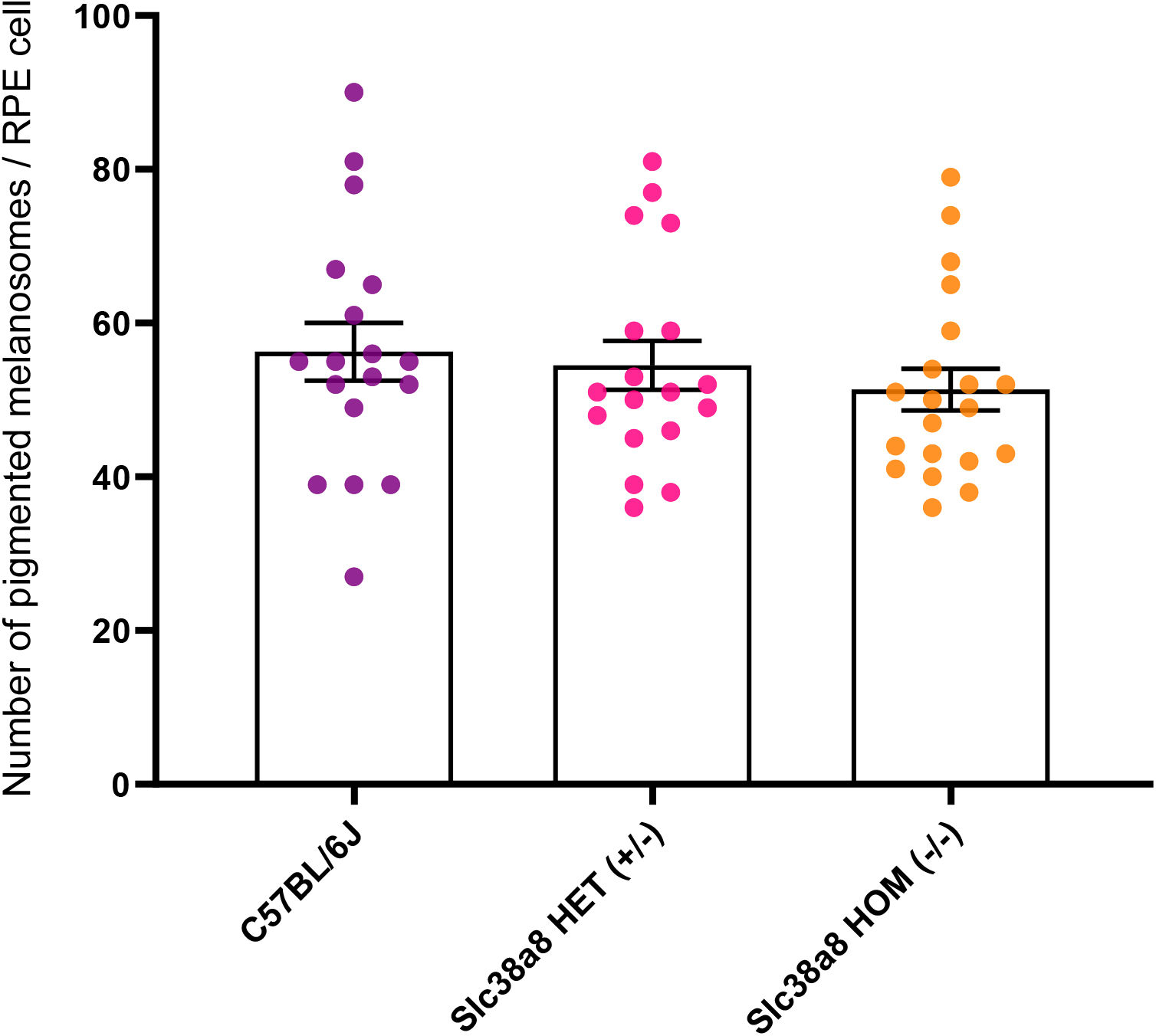
Number of pigmented melanosomes per RPE cell, counted from electron microscope images, in adult (2-3 months old) wild-type pigmented C57BL/6J, FHONDA *Slc38a8* heterozygous “HET” (+/−) and FHONDA *Slc38a8* homozygous mutant “HOM” (−/−). Mean ± SEM, N=18-20 RPE cells, 1-factor ANOVA with Bonferroni correction for multiple comparisons. No statistically significant differences were detected.

